# Mild replication stress causes premature centriole disengagement via a sub-critical Plk1 activity under the control of ATR-Chk1

**DOI:** 10.1101/2022.09.29.510042

**Authors:** Devashish Dwivedi, Daniela Harry, Patrick Meraldi

## Abstract

A tight synchrony between the DNA and centrosome cycle is essential for genomic integrity. Centriole disengagement, which licenses centrosomes for duplication, occurs normally during mitotic exit. We recently demonstrated that mild DNA replication stress in untransformed human cells causes premature centriole disengagement at mitotic entry, leading to transient multipolar spindles that favour chromosome mis-segregation. How mild replication stress accelerate the centrosome cycle at the molecular level remained, however, unclear. Using expansion microscopy, we show that mild replication stress already induces premature centriole disengagement in G2 via the ATR-Chk1 axis of the DNA damage repair pathway. We demonstrate that this results in a subcritical Plk1 kinase activity that is insufficient for rapid mitotic entry. Nevertheless, it primes the pericentriolar matrix for Separase-dependent disassembly causing premature centriole disengagement in G2. We postulate that the differential requirement of Plk1 activity in the DNA and centrosome cycles explains how mild replication stress disrupts the synchrony between both processes and contributes to genomic instability.

## Introduction

The centrosomes in animal cells are the major microtubule organising centre (MTOC) that regulate the interphase microtubule network and control the poles of the mitotic spindle during cell division. Centrosomes also integrate and coordinate multiple signalling pathways involved in regulation of cell polarity, migration, development, and fate^1^. Centrosomal dysfunctions may result in developmental disorders or cancer^2-4^. Each centrosome consists of two tightly associated and orthogonally oriented barrel-shaped centrioles that are 500 nm long and 250 nm wide, called mother and daughter centrioles. These two centrioles are surrounded with a protein rich pericentriolar matrix (PCM). In dividing animal cells, the two centrioles within centrosomes duplicate once per cell cycle in a process that is tightly controlled in space and time^5^. The centriole duplication cycle starts with centriole disengagement during telophase of the previous cell cycle. Indeed, a steric blockade inhibits the formation of new procentrioles as long as both centrioles within the centrosome are in tight orthogonal association with each other^6, 7^. Centriole disengagement is under the control of the mitotic kinase Plk1 and the protease Separase^8, 9^, with Plk1 targeting by phosphorylating the pericentriolar protein pericentrin for Separase-dependent cleavage^8-11^. As centrioles disengage, they lose their orthogonal orientation and steric hindrance, licensing them for duplication during the next S-phase^12^. The duplicated centrosomes, each containing a pair of engaged centrioles, separate at mitotic onset to form the poles of the bipolar spindle, before being segregated into the two daughter cells during anaphase.

Under normal conditions the centriole duplication cycle proceeds synchronously with the cell cycle as both DNA and centrosomes are licensed for replication at the end of mitosis, replicated in S-phase and segregated to the two daughter cells during mitosis^5^. Consistently, both processes share multiple common regulatory proteins^13^. Any dysregulation in the centriole duplication cycle can lead to the formation of abnormal spindles in mitosis, in particular multipolar spindles which are commonly observed in many cancers^14^. Even tough cancers cells possess centrosome clustering mechanisms to convert multipolar spindles into bipolar spindles, transient abnormal spindles will favour erroneous chromosome attachments to spindle microtubules, increasing the probability of chromosome instability (CIN) and aneuploidy^15, 16^

We recently reported that mild replication stress induced by nanomolar doses of the DNA-polymerase inhibitor Aphidicolin causes premature centriole disengagement in mitosis, disrupting the synchrony between the centrosome- and the cell-cycle. Premature centriole disengagement in cells under replication stress resulted in transient multipolar spindles that often led to chromosome segregation errors and chromosome instability in anaphase^17^. Replication stress is recognised as any cellular condition in which DNA replication is slowed down or hampered, a condition which is already present in many pre-cancerous lesions^18^. It can result in the formation of double or single DNA strand breaks (DSBs/SSBs), which activate the damage repair (DDR) kinases Ataxia telangiectasia mutated (ATM; in case of DSBs) or Ataxia Telangiectasia and Rad3-related protein (ATR; in case of SSBs) to delay mitotic onset^19^. The link between the DDR pathway and the centriole duplication cycle, however, remains so far unclear. Here, we combined small molecule-based inhibition against different cell cycle and DDR regulators and protein depletions with ultrastructure expansion microscopy (UxM) to unravel the molecular signalling pathway involved in premature centriole disengagement under mild replication stress conditions. Our results show that centriole disengagement depends on the ATR-Chk1 axis of DDR pathway and that it is caused by creating sub-critical level of Plk1 activity that suffice to drive premature centriole disengagement via the canonical Plk1-Separase-pericentrin cleavage mechanism in G2, but are insufficient to promote efficient mitotic entry itself.

## Results

### Mild replication stress induces premature centriole disengagement in G2

To investigate how mild replication stress disrupts the synchrony between the cell- and centriole duplication-cycle, we worked with untransformed human retinal pigment epithelial (hTERT-RPE1) cells immortalised with human telomerase. These cells have functional cell-cycle checkpoints, low basal incidences of chromosome segregation errors and normal centriole duplication cycle. To induce mild replication stress, we applied low doses (400 nM) of the DNA polymerase Aphidicolin for 16 hrs, a condition known to induce premature centriole disengagement in mitosis^17^. To study the origin of premature centriole disengagement we used ultrastructure expansion microscopy (UxM) to test whether low doses of Aphidicolin already induced centriole disengagement in S- or G2-phase. This methodology allowed to overcome the resolution limit of conventional light microscopy and to determine whether the centriole pair within a centrosome was engaged or not (Fig. 1 A & B). Centrioles that were closely associated (<500 nm apart i.e., one centriole length distance) and in perfect orthogonal orientation were considered engaged; centriole pairs that did not meet these criteria were classified as disengaged (Fig. 1A and B).

**Figure 1:**
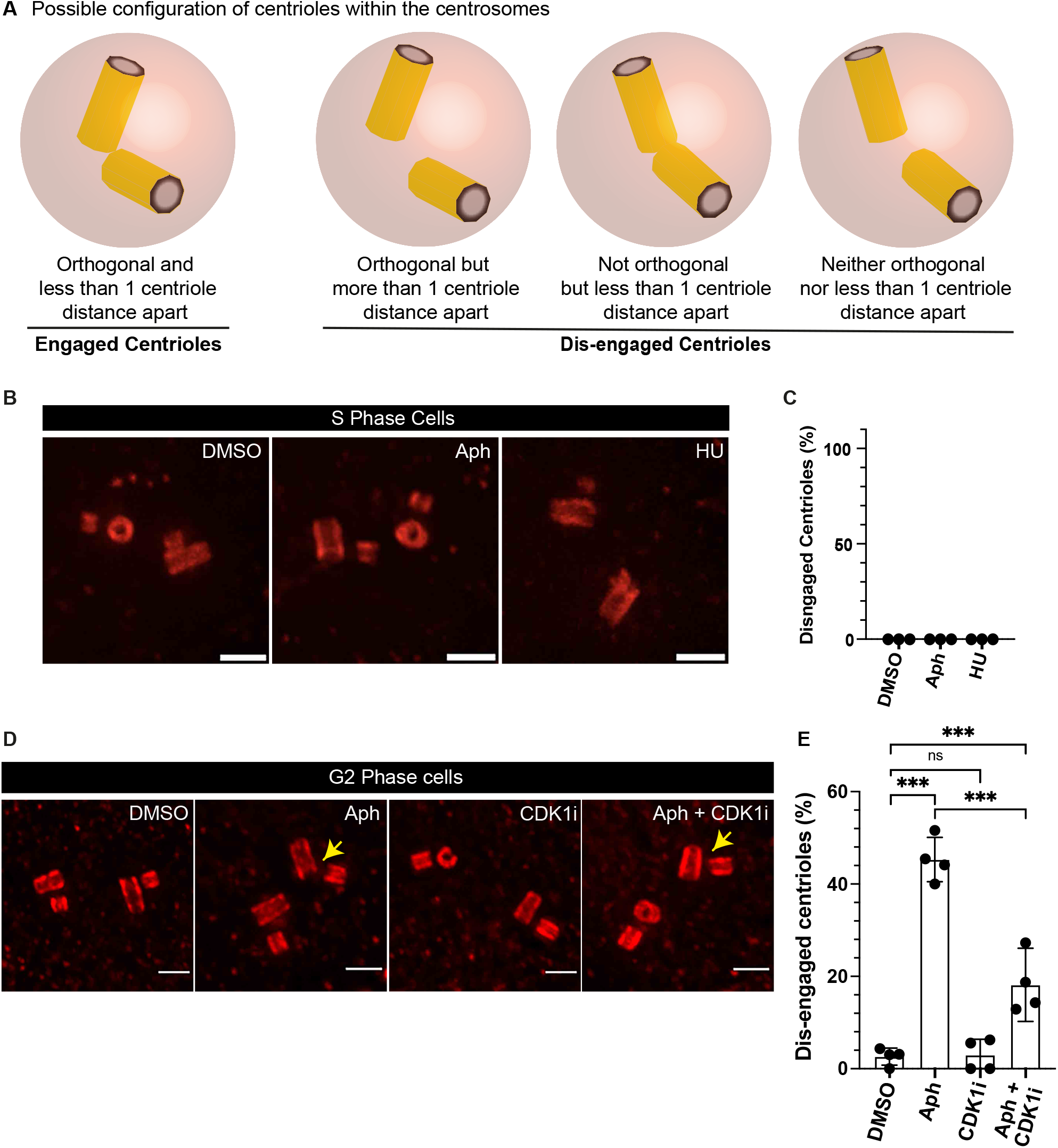
Mild replication stress causes premature centriole disengagement in G2 phase. **(A)** Possible centriole configurations in G2 centrosomes to identify centrosomes with disengaged centrioles. **(B)** Expansion microscopy images of centrioles in S phase hTERT-RPE1 cells stained against α-tubulin and treated with indicated drugs. **(C)** Quantification of percentage of S phase cells with dis-engaged centrioles in their centrosomes (*N* = 3 independent experiments, n = 86, 79 and 81 cells for DMSO, Aph and HU respectively). **(D)** Expansion microscopy images of centrioles in G2 phase hTERT-RPE1 cells stained against and α-tubulin treated with indicated drugs/inhibitors. **(E)** Quantification of percentage of G2 phase cells with dis-engaged centrioles in their centrosomes (*N* = 4 independent experiments, n = 111, 128, 101 and 111 cells in DMSO, Aph, Cdk1i and Aph+Cdk1i respectively). Data presented as Mean values ± SD. (p > 0.05 = ns; p < 0.001 = ***: Sídak test). Scale bars = 0.5 µm.

In S-phase cells, which contained short pro-centrioles that were not fully elongated (<50% length of the parental centriole), all centriole pairs were engaged whether cells were treated with DMSO (negative control) or 400 nM Aphidicolin (Fig. 1B & C). In contrast, in G2 cells treatment with 400 nM Aphidicolin led to centriole disengagement in 45.30±4.81% of the cells vs 2.63±1.85% in DMSO-treated cells (Fig. 1D-E). Our previous study indicated that Cdk1 activity was required during the low Aphidicolin treatment to induce premature centriole disengagement in mitosis^17^. Consistent with these findings, we found that Cdk1 inhibition in Aphidicolin treated cells using the small molecule inhibitor RO-3306^20^ also suppressed centriole disengagement in G2 cells (18.14±6.47%; Fig. 1D-E) when compared to aphidicolin treated cells. Cdk1 inhibition on its own did not affect centriole engagement in G2 cells (2.95±3.42%; Fig. 1D). We conclude that mild replication stress already induces centriole disengagement in G2 cells, via the same mechanisms leading to premature centriole disengagement in mitosis.

### ATR-Chk1 regulate premature centriole disengagement under mild replication stress conditions

Replication stress induces activation of the DDR pathway^19^ and delays mitotic entry by engaging the G2/M checkpoint^21^ (Fig. 2A). We therefore mapped out which part of the DNA damage machinery might regulate centriole disengagement in G2 under mild replication stress conditions. We first inhibited the upstream DNA damage repair kinases ATR (Ataxia telangiectasia and Rad3-related) and ATM (ataxia-telangiectasia mutated) in Aphidicolin treated cells using the selective small molecular inhibitors ETP46464 (ATR inhibitor) and KU55933 (ATM inhibitor)^22, 23^. Inhibiting ATR activity suppressed the Aphidicolin-induced centriole disengagement in G2 phase (7.40±3.70% vs. 45.42±3.03% with Aphidicolin alone; Fig. 2B-C). Inhibiting ATM activity under similar replication stress conditions did not prevent premature centriole disengagement in G2 (40.96±3.93%; Fig 2B-C). Finally, treatment with ATM (10.09±1.26%) or ATR (11.76±9.89%) inhibitors alone led to a very mild increase in centriole disengagement (3.13±0.08%). We conclude that premature centriole disengagement in G2 under mild replication stress conditions depends on ATR but not on ATM kinase activity.

**Figure 2:**
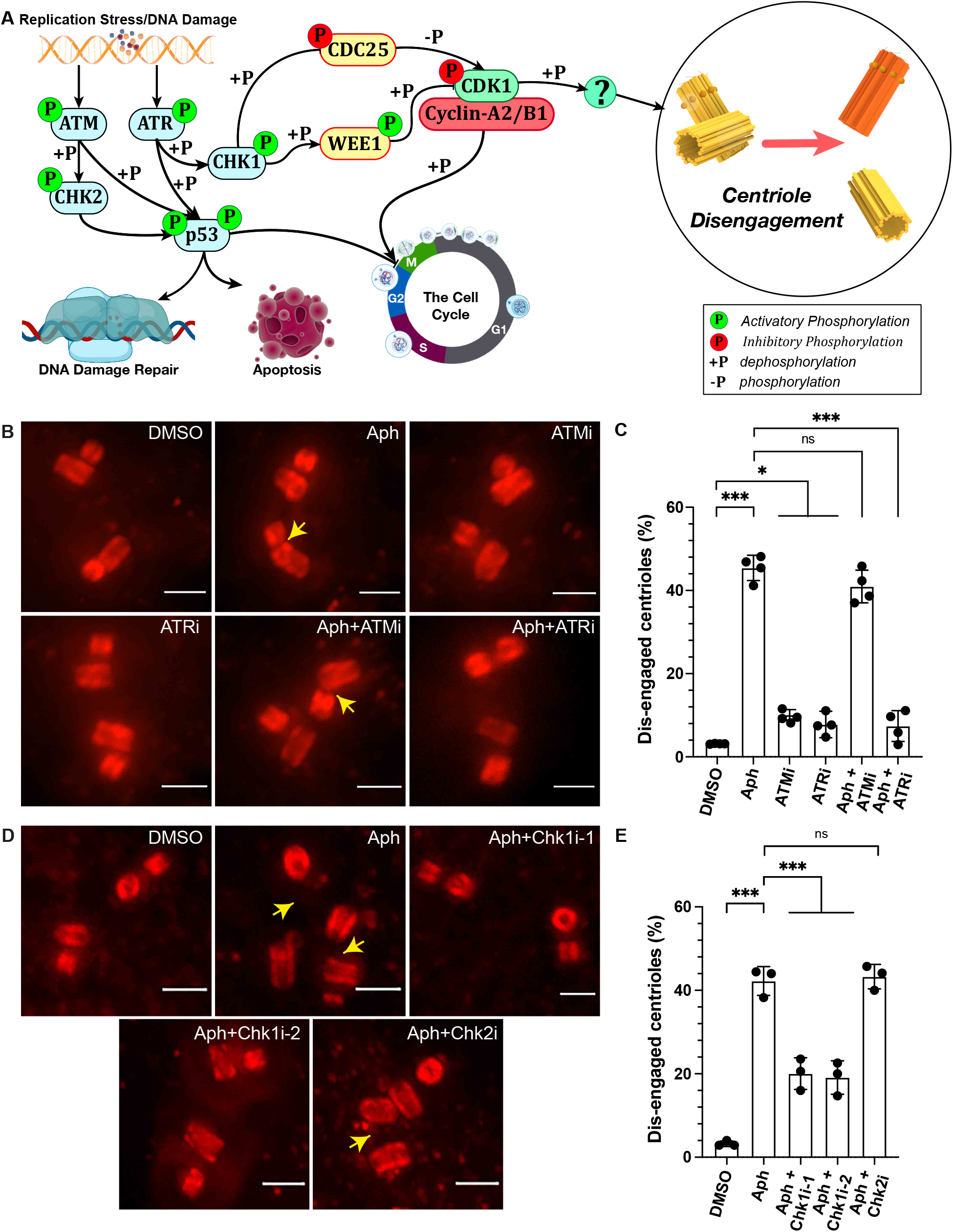
Premature centriole disengagement during mild replication stress depends on ATR/Chk1 axis of DDR Pathway. **(A)** Current model for pathways linking DNA damage and cell cycle control. **(B)** Expansion microscopy images of centrioles in G2 Phase hTERT-RPE1 cells, treated with indicated drugs/inhibitors targeting the DNA damage repair pathway. **(C)** Quantification of percentage of G2 phase cells with engaged centrioles in their centrosomes (*N* = 3 independent experiments, n = 96, 93, 70, 71, 76 and 92 cells in DMSO, Aph, ATMi, ATRi, Aph+ATMi and Aph+ATRi respectively). **(D)** Expansion microscopy images of centrioles in G2 Phase hTERT-RPE1 cells, treated with indicated drugs/inhibitors targeting Chk1 or Chk2. **(E)** Quantification of percentage of G2 phase cells with engaged centrioles in their centrosomes (*N* = 3 independent experiments, n = 95, 95, 92, 90 and 94 cells in DMSO, Aph, Aph+Chk1i-1, Aph+Chk1i-2 and Aph+Chk2i respectively). Data presented as Mean values ± SD. (p > 0.05 = ns; p < 0.001 = ***: Sídak test). Scale bars = 0.5 µm.

To initiate DNA repair and prevent mitotic entry in cells having damaged DNA, ATR activates the protein kinase Chkl, while ATM activates the Chk2 kinase (Fig. 2A). We therefore next suppressed the activity of Chk1 with two independent inhibitors, LY2603618 (Chk1i-1) and PF-477736 (Chkk1i-2)^24, 25^ and Chk2 using small molecule inhibitors (Chk2i)^26^ in Aphidicolin treated cells and monitored the centriole engagement status. None of the inhibitors affected centriole engagement when administered alone (3.85±0.68%: Chk1i-1, 5.21±2.60%: Chk1i-2, 5.07±2.70%: Chk2; Fig. S1A-B). Chk1 inhibition, however, suppressed premature centriole disengagement in G2 in Aphidicolin-treated cells (20.04±3.78%: Chk1i-1 and 19.09±4.01%: Chk1i-2), whereas Chk2 inhibition had no effect (43.28±2.95% vs 42.22±3.46% in Aphidicolin-treatment alone; Fig. 2D-E). We conclude that the premature centriole disengagement in G2 induced by mild replication stress depends on the ATR-CHK1 arm of the DNA damage repair pathway.

### Premature centriole disengagement under replication stress depends on Wee1 and Cdk1/Cyclin-A

The ATR-Chk1 arm of the DNA damage repair pathway activates the Wee1 kinase which phosphorylates Cdk1 at Tyrosine-15 to prevent mitotic entry and prolongs G2^27, 28^. To evaluate to which extend Wee1 activity and/or a prolonged G2 phase is required for a premature centriole disengagement, we inhibited Wee1 using the small molecule inhibitors PD0166285 (Wee1i-1) and MK1775 (wee1i-2)^29, 30^. Since such a treatment dramatically shortens G2, we could not analyse the centriole configuration in G2 cells by UxM. Instead, we used live-cell imaging to detect transient multipolar mitotic spindles that arise after Aphidicolin treatment^17^. While RPE1 cells expressing EB3-GFP (mitotic spindle marker) and H2B-mCherry (DNA marker) underwent normal cell divisions after DMSO treatment (0.52±0.57% multipolar spindles - Movie 1), Aphidicolin treatment significantly favoured the formation of transient multipolar spindles during early mitosis (15.45±1.19%; Fig. 3A-B - Movie 2), as previously observed^17^. Inhibition of Wee1 on its own did not change the frequency of transient multipolar spindle (0.66±0.11%: Wee1i-1 and 0.80±0.16%: Wee1i-2 - Movie 3 & 4), but suppressed this phenotype in Aphidicolin-treated cells (4.49±0.59%: Wee1i-1 and 4.38±0.41%: Wee1i-2; Fig. 3A-B - Movie 5 & 6). We conclude that Wee1 activity is required for premature centriole disengagement after mild replication stress.

**Figure 3:**
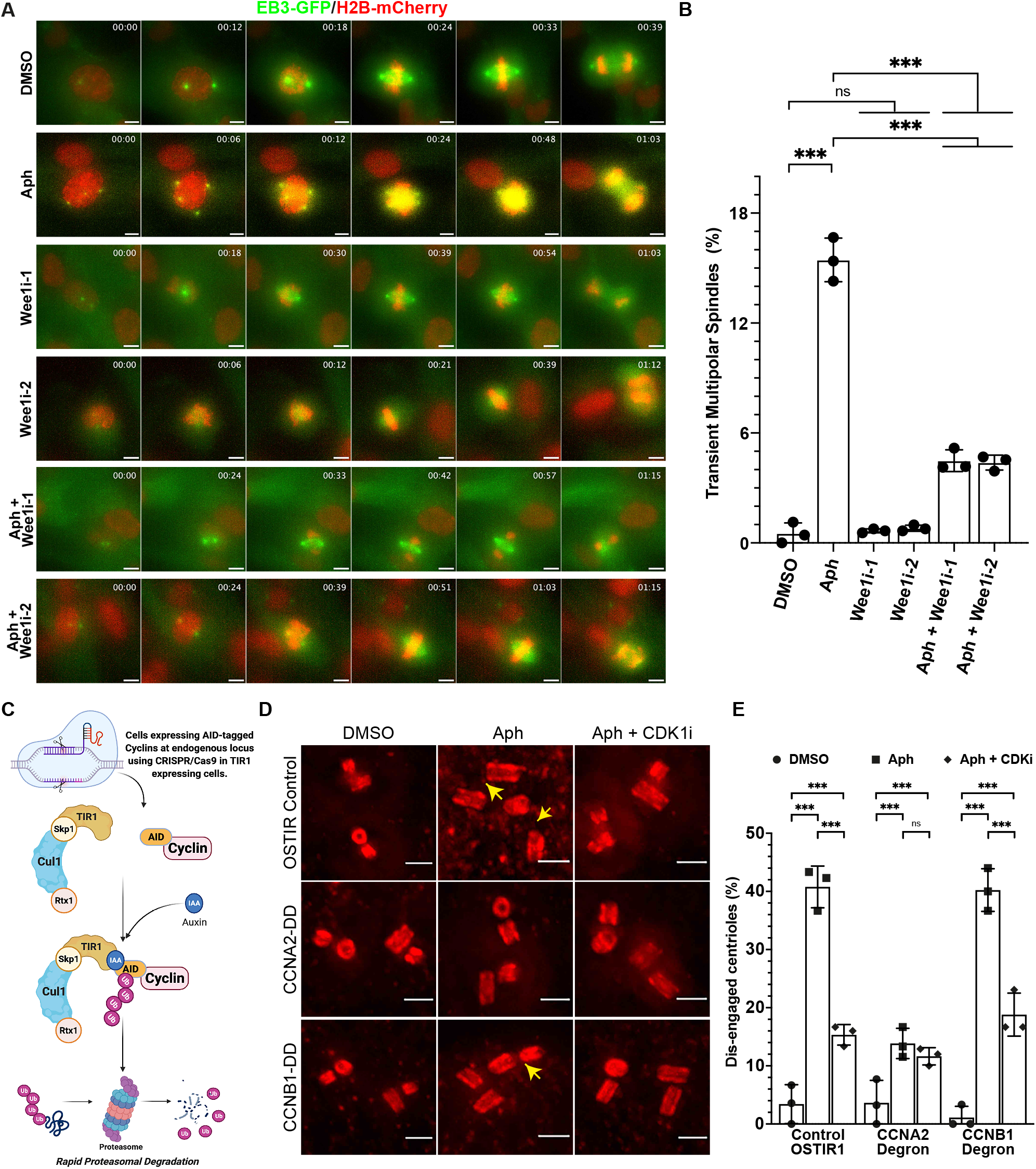
Premature centriole disengagement depends of Wee1 kinase and Cdk1/Cyclin-A2. **(A)** Live-cell time-lapse images of hTERT-RPE1 EB3-eGFP/H2B-mCherry cells treated with indicated drugs/inhibitors. Scale bars = 5 µm. **(B)** Quantification for the proportion of mitotic cells displaying transient multipolar spindles upon mitotic entry in (A) (*N* = 3 independent experiments, n = 485, 137, 462, 385, 333 and 276 cells in DMSO, Aph, Wee1i-1, Wee1i-2, Aph+Wee1i-1 and Aph+wee1i-2, respectively). **(C)** Schematic representation of auxin induced cyclin degradation of Cyclin-A2 and B1 in hTERT-RPE1 cells. **(D)** Expansion microscopy images of centrioles in G2 Phase hTERT-RPE1 OSTIR1, Cyclin-A2 double degron (CCNA2-DD) and Cyclin-B1 double degron (CCNB2-DD) cells after inducing degron expression and treated with indicated drugs/inhibitors. **(E)** Quantification of percentage of G2 phase cells with dis-engaged centrioles in their centrosomes [*N* = 3 independent experiments, n = 88 (OSTIR:DMSO), 86 (OSTIR:Aph), 85 (OSTIR:Aph+Cdk1i), 88 (CCNA2-DD:DMSO), 86 (CCNA2-DD:Aph), 86 (CCNA2-DD:Aph+Cdk1i), 85 (CCNB1-DD:DMSO), 75 (CCNB1-DD:Aph) and 70 (CCNB1-DD:Aph+Cdk1i) cells]. Data presented as Mean values ± SD. Data presented as Mean values ± SD. (p > 0.05 = ns; p < 0.001 = ***: Sídak test). Scale bars = 0.5 µm.

Even though Cdk1 inhibition is the main action of Wee1^28^, premature centriole disengagement requires Cdk1 activity^17^. This activity is driven in late G2 phase by two cyclins: Cyclin-A2 and Cyclin-B1 which are expressed at similar levels at this stage. To determine which cyclin drives centriole disengagement in G2, we used established RPE1 cells expressing auxin inducible degron (AID) endogenously tagged Cyclin-A2 or Cyclin-B1^31^, which allow rapid degradation of the tagged protein by the ubiquitin/proteasome system (Fig 3C and validation of depletion in Fig. S2A-B). Degradation of Cyclin-A2 or Cyclin-B1 alone led to normal rates of premature centriole disengagement in G2 when compared to the parental cell line expressing only the E3-ubiquitin ligase OSTIR (3.41±3.34%: OSTIR; 3.64±3.86%: Cyclin-A2-AID and 1.11±1.92%: Cyclin-B1-AID; Fig. 3D-E). However, premature centriole disengagement induced by Aphidicolin in G2 was suppressed upon Cyclin-A2 degradation, but not by Cyclin-B1 degradation (13.85±2.60%: Cyclin-A2, Cyclin-B1: 40.22±3.67%, 40.77±3.59%: OSTIR; Fig. 3D-E). In line with a Cdk1/CyclinA-2 dependent centriole disengagement, Cdk1 inhibition only suppressed premature centriole disengagement in control- (15.33±1.77%) and Cyclin-B1-AID (18.81±3.70) cells, but had no further effect in Cyclin-A2-AID cells (11.63±1.48%; Fig. 3C-D). We conclude that premature centriole disengagement in G2 is driven by Cdk1/CyclinA-2.

### Replication stress results in intermediate Plk1 activity

One of the prominent downstream effectors of Cdk1/Cyclin-A is the protein kinase Plk1^32, 33^. Plk1 is required for the physiological centriole disengagement^11, 34, 35^ in telophase and the premature centriole disengagement in mitosis after replication stress^17^. Moreover, Plk1 overexpression can induce centriole disengagement^34^. We therefore inhibited Plk1 in Aphidicolin treated cells using small molecule inhibitor, BI-2536^36^ and quantified centriole disengagement in G2. We found a strong suppression of centriole disengagement (13.41±0.77%) when compared to cells treated with Aphidicolin alone (46.04±0.18%), and no significant change in the absence of Aphidicolin (3.77±0.37%: Plk1 inhibition vs 2.62±2.27%: DMSO; Fig. 4A-B). We conclude that premature centriole disengagement in G2 also depends on Plk1 activity.

**Figure 4:**
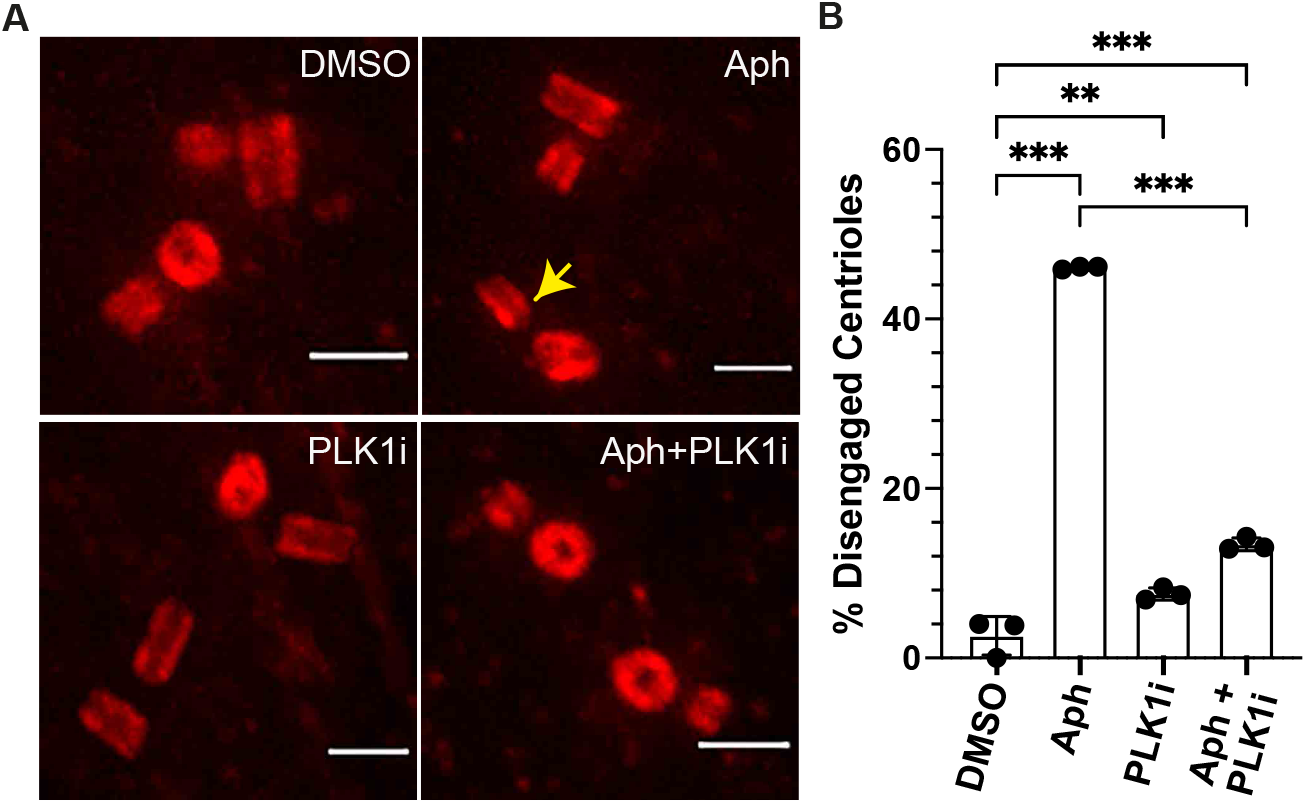
Premature centriole disengagement requires Plk1 activity. **(A)** Expansion microscopy images of centrioles in G2 phase hTERT-RPE1 cells treated with indicated drugs/inhibitors. **(B)** Quantification of percentage of G2 phase cells with engaged centrioles in their centrosomes (*N* = 3 independent experiments, n = 79, 76, 80 and 82 cells in DMSO, Aph, Plk1i, and Aph+Plk1i respectively). Data presented as Mean values ± SD. (p < 0.01 = **; p < 0.001 = ***: Sídak test). Scale bars = 0.5 µm.

Overall, our results pointed to a paradox: premature centriole disengagement in G2 depends on the ATR-Chk1-Wee1 pathway that limits Cdk1 and Plk1 activity, yet at the same time it depended on Cdk1/CyclinA2 and Plk1. We therefore hypothesized that mild replication stress might lead to a partial Plk1 activity that would be sufficient to drive centriole disengagement, but not sufficient to rapidly enter mitosis. To test this hypothesis, we quantified Plk1 activity using a Föster’s Resonance Energy Transfer (FRET) Plk1 activity sensor, based on a c-Jun peptide (Plk1 substrate) coupled to CFP and YFP^37^. Phosphorylation of this peptide induces a confirmational change that decreases the FRET efficiency allowing to quantify Plk1 activity. Quantitative immunofluorescence of RPE1 cells expressing the Plk1-FRET sensor indicated high FRET (YFP to CFP ratio) values in cells arrested in G1 after Cdk4/6 inhibition (no Plk1 activity; 3.50±0.50), in G2 cells treated with a Cdk1 inhibitor (no Plk1 activation; 3.79±0.45) or in prometaphase cells treated with a Plk1 inhibitor (3.75±0.45), but low FRET values (1.20±0.45) in cells arrested in prometaphase with a Eg5^KIF11^ inhibitor (fully active Plk1; Fig 5A-B). Using these values as internal standards, we found that low doses of Aphidicolin decreased Plk1 activity in a dose dependent manner: 200 nM (1.91±0.0.54), 400 nM (2.35±0.66), 600 nM (3.65±0.58) and 800 nM (3.77±0.57; Fig 5A-B). This indicated that mild replication stress (400nM Aphidicolin) resulted in an intermediate (45% of maximal) Plk1 activity. This intermediate Plk1 activity was suppressed by Cdk1 or Plk1 inhibition (Aph+Cdk1i: 3.39±0.58; Aph+Plk1i: 3.53±0.53; Fig. 5C-D). We conclude that mild replication stress leads to an intermediate Cdk1-dependent Plk1 activity in G2.

**Figure 5:**
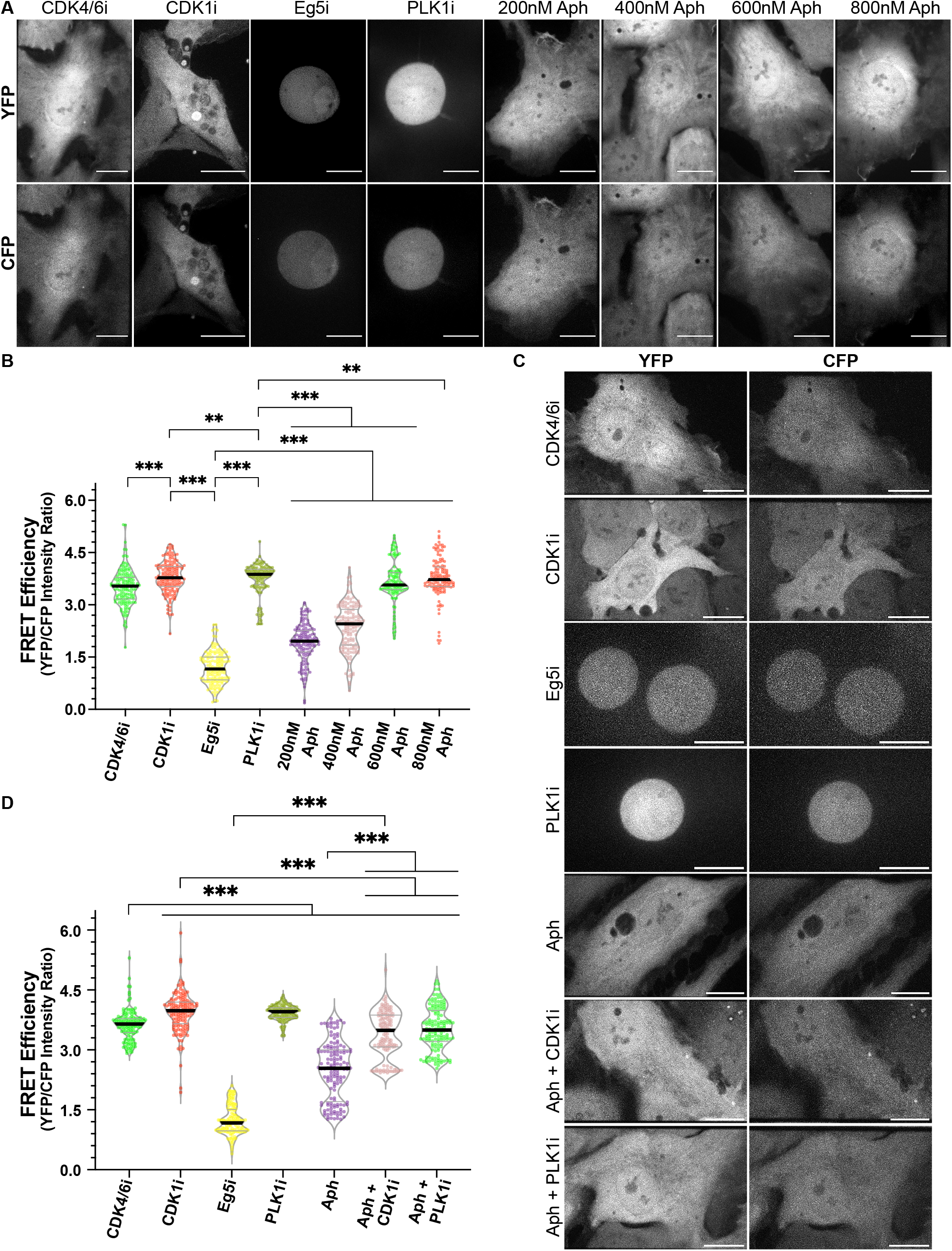
Mild replication stress induces sub-critical Plk1 activity. **(A)** Representative images of CFP and YFP fluorescence from hTERT-RPE1 Plk1-FRET Sensor cells treated with indicated inhibitors/drugs. **(B)** Violin plots of YFP/CFP intensity ratios from cells in A [*N* = 3 independent experiments, n = 162 (Cdk4/6i), 164 (Cdk1i), 157 (Eg5i), 164 (Plk1i), 161 (200 nM Aph), 167 (400 nM Aph), 134 (600 nM Aph) and 132 (800 nM Aph) cells]. **(C)** Representative images of CFP and YFP fluorescence from hTERT-RPE1 Plk1-FRET Sensor cells treated with indicated inhibitors/drugs. **(D)** Violin plots of YFP/CFP intensity ratios from cells in C [*N* = 3 independent experiments, n = 149 (Cdk4/6i), 149 (Cdk1i), 142 (Eg5i), 154 (Plk1i), 148 (Aph), 135 (Aph+Cdk1i) and 143 (Aph+Plk1i)]. Median in each case is marked with bold black line and thin grey lines denote the 1^st^ and 3^rd^ quartiles. (p < 0.01 = **; p < 0.001 = ***: Sídak test). Scale bars = 0.5 µm.

### ATR-Chk1 impose an intermediate Plk1 activity under mild replication stress

Activation of multiple players in the DDR pathway require Plk1 activity and Plk1 controls the recovery from G2/M arrest during DNA damage once the DNA repair is completed to facilitate mitotic entry ^38-40^. To test whether the intermediate Plk1 activity is controlled by the ATR-Chk1 axis in Aphidicolin-treated cells we again quantified Plk1 activity using the FRET based reporter. Inhibition of ATR or Chk1 kinase activity in Aphidicolin treated cells fully rescued Plk1 activity as indicated by low FRET ratio values (1.20±0.16: ATRi, 1.29±0.09: Chki vs. 2.28±0.66 Aphidicolin alone; Fig. 6A & B). In contrast, ATM or Chk2 inhibition had no effect on Plk1 activity in Aphidicolin treated cells (2.50±0.29: ATMi, 2.24±0.49: Chk2i; Fig. 6A-B). This effect was specific for Aphidicolin-treated cells, since ATR or Chk1 inhibition had no effect on Plk1 activity in prometaphase arrested cells with an Eg5 inhibitor (1.20±0.16: Eg5i+ATRi, 1.13±0.27: Eg5i+Chk1i vs. 1.18±0.25: Eg5i). We conclude that under mild replication stress conditions the ATR-Chk1 axis but not the ATM-Chk2 axis lowers Plk1 activity to an intermediate level, consistent with studies indicated that Chk1 acts upstream of Plk1 and that Chk1 can directly phosphorylate Plk1^41, 42^.

**Figure 6:**
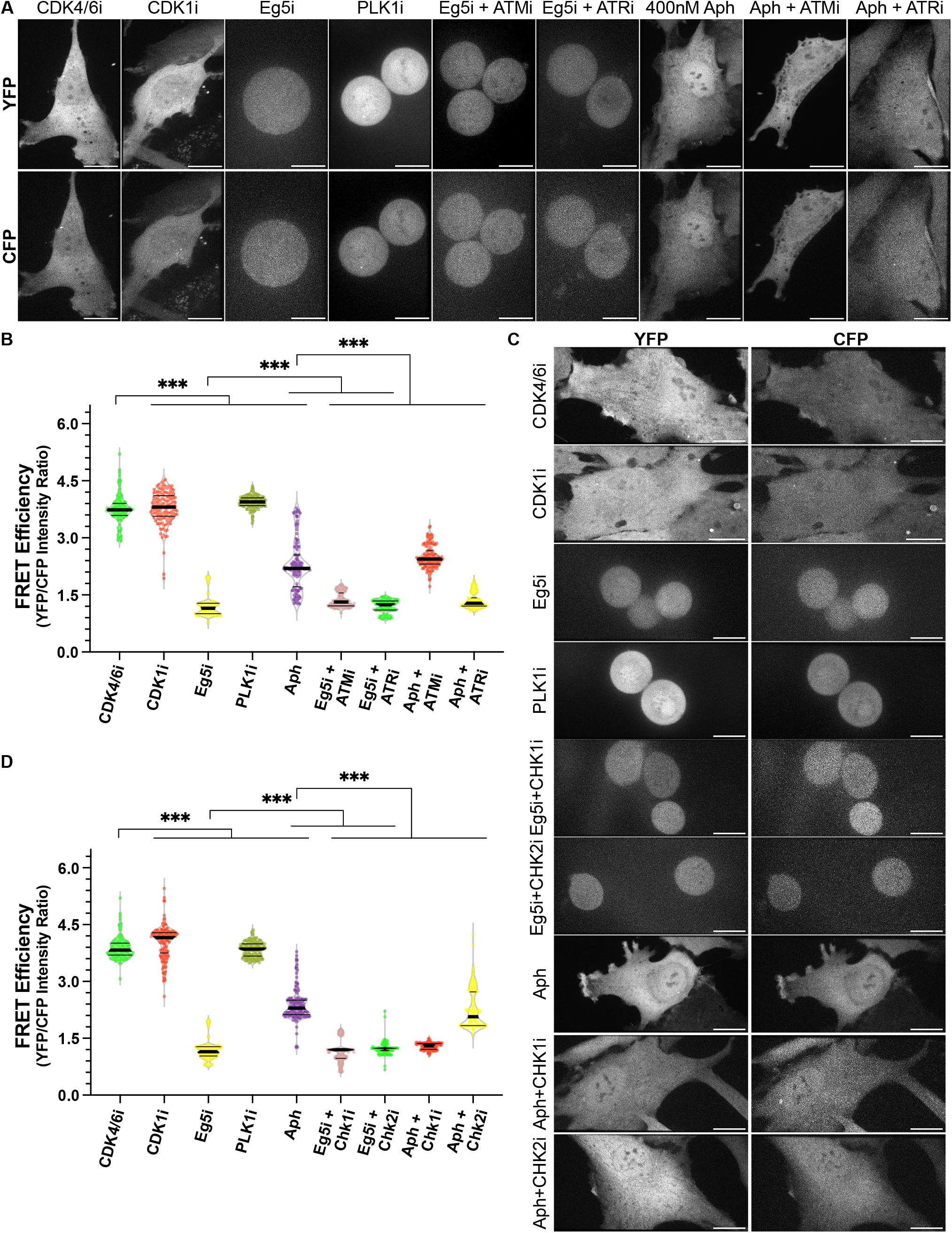
ATR/Chk1 axis regulates the sub-critical Plk1 activity of during replication stress. **(A)** Representative images of CFP and YFP fluorescence from hTERT-RPE1 Plk1-FRET Sensor cells treated with indicated inhibitors/drugs targeting indicated actors of the DNA damage repair pathway. **(B)** Violin plots of YFP/CFP intensity ratios from cells in A [*N* = 3 independent experiments, n = 146 (Cdk4/6i), 146 (Cdk1i), 146 (Eg5i), 151 (Plk1i), 149 (Aph), 151 (Eg5i+ATMi), 156 (Eg5i+ATRi), 159 (Aph+ATMi) and 159 (Aph+ATRi)]. **(C)** Representative images of CFP and YFP fluorescence from hTERT-RPE1 Plk1-FRET Sensor cells treated with indicated inhibitors/drugs against DNA repair pathway proteins. **(D)** Violin plots of YFP/CFP intensity ratios from cells in C [*N* = 3 independent experiments, n = 152 (Cdk4/6i), 173 (Cdk1i), 178 (Eg5i), 175 (Plk1i), 176 (Aph), 180 (Eg5i+Chk1i), 181 (Eg5i+Chk2i), 176 (Aph+Chk1i) and 185 (Aph+Chk2i)]. Median in each case is marked with bold black line and thin grey lines denote the 1^st^ and 3^rd^ quartiles. (p < 0.001 = ***: Sídak test). Scale bars = 0.5 µm.

### Plk1 activates Separase and primes pericentrin to promote centriole disengagement

Plk1 induces centriole disengagement during mitotic exit by priming the pericentriolar protein Pericentrin for localized cleavage by the protease Separase ^8-10^. To test whether Plk1 promotes premature centriole disengagement via the same molecular pathway, we studied the localization of pericentrin in G2 cells with or without replication stress by super-resolution Stimulated emission depletion (STED) microscopy. While in DMSO-treated G2 cells 95.48±1.38% of the centrioles displayed a ∼350 nm pericentrin ring, only 15.71±3.56% of the centrioles in Aphidicolin-treated cells displayed such a complete ring (Fig. 7A-B and S3A-B). An equivalent disruption of the Pericentrin ring was also found in control telophase cells (DMSO) when centrioles disengage normally (Fig.7A and S3C). Finally, a very similar loss of the pericentrin ring integrity could be observed after Aphidicolin treatment when using UxM (Fig. 7C-D). In contrast, the localization of CEP57, another centriolar protein responsible for maintaining centriole engagement during mitosis ^43^ showed no change in localization after Aphidicolin treatment (Fig. 7E-F).

**Figure 7:**
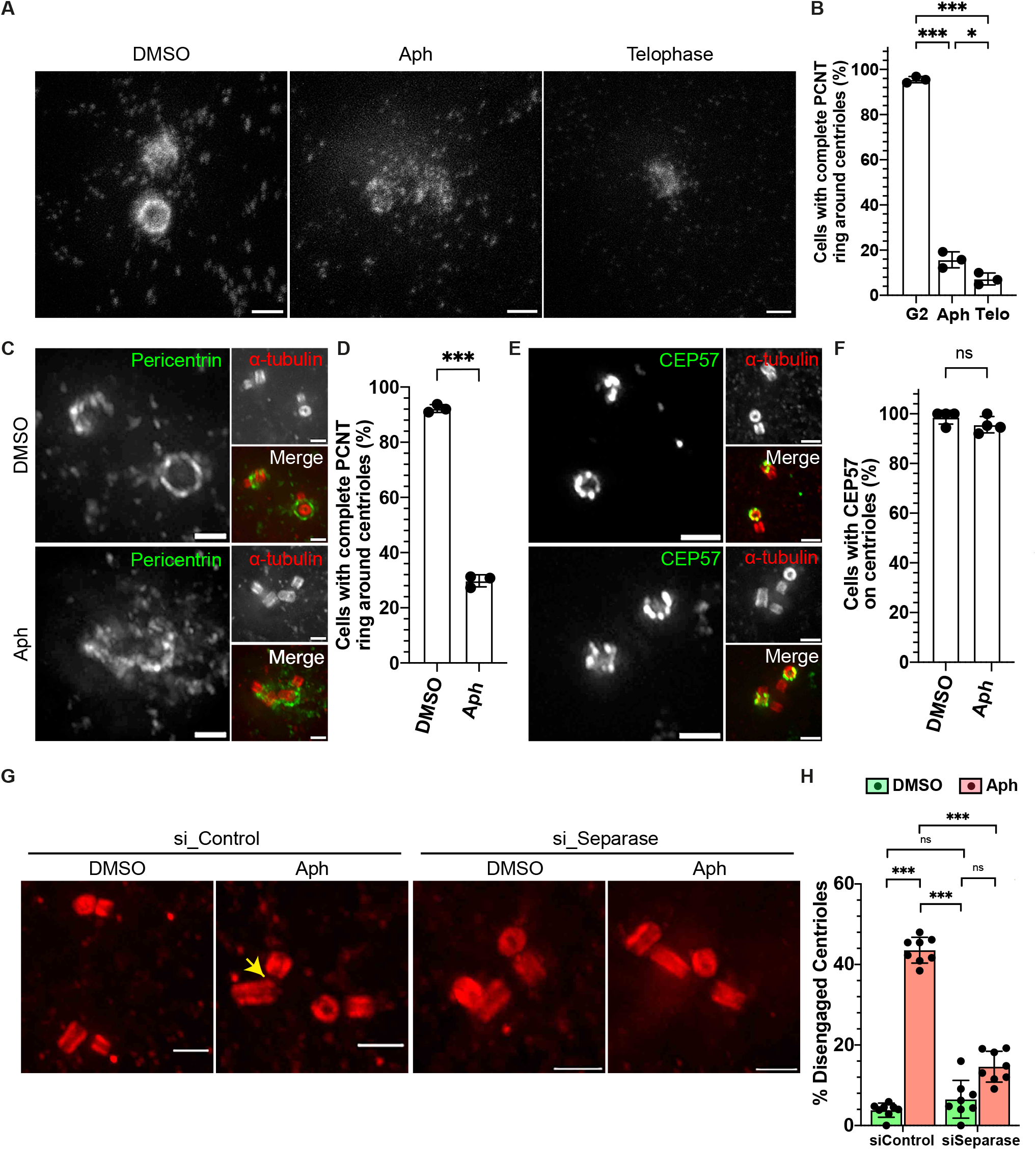
Mild replication stress affects PCM integrity. **(A)** STED nanoscopy images of zoomed pericentriolar region of G2 phase hTERT-RPE1 cells treated either with DMSO or Aph and DMSO treated cells in telophase. **(B)** Quantification for the percentage of G2 cells having intact pericentrin as ring in conditions depicted in A (*N* = 3 independent experiments, n = 71, 78 and 70 for DMSO: G2 phase, Aph and DMSO: Telophase cells, respectively). **(C)** Expansion microscopy images of centrioles in G2 Phase hTERT-RPE1 cells treated either with DMSO or Aph and stained for α-tubulin (red) and Pericentrin (green). **(D)** Quantification of percentage of G2 phase cells with complete Pericentrin ring around their centrioles (*N* = 4 independent experiments, n = 82 and 76 cells in DMSO and Aph, respectively). **(E)** Expansion microscopy images of centrioles in G2 Phase hTERT-RPE1 cells treated either with DMSO or Aph and stained for α-tubulin (red) and CEP57 (green). **(F)** Quantification of percentage of G2 phase cells with complete CEP57 ring around their centrioles (*N* = 4 independent experiments, n = 65 and 83 cells in DMSO and Aph, respectively). **(G)** Expansion microscopy image of centrioles in G2 phase hTERT-RPE1 cells treated either with siControl or siSeparase and DMSO or Aph. **(H)** Quantification of percentage of G2 phase cells having engaged centrioles (*N* = 8 independent experiments, n = 187, 195, 188 and 192 cells in siControl:DMSO, siControl:Aph, siSeparase:DMSO and siSeparase:Aph, respectively). Data presented as Mean values ± SD. (p > 0.05 = ns; p < 0.05 = *; p < 0.001 = ***: Sídak test). Scale bars = 0.5 µm.

Next, we quantified the contribution of Separase to premature centriole disengagement. We depleted Separase by siRNA in Aphidicolin or DMSO-treated cells (Fig. 7G) and monitored centriole disengagement. While Separase depletion had no effect on centriole disengagement in the absence of replication stress (Control siRNA: 3.81±1.76%; Separase siRNA: 6.52±4.70%; Fig. 7G-H), it strongly suppressed premature centriole disengagement in G2 in cells treated with Aphidicolin (14.62±3.84% vs. 43.55±3.20% in *siControl;* Fig. 7G-H). Moreover, Separase depletion restored in large parts the Pericentrin ring integrity around centrioles in Aphidicolin treated cells G2 cells (76.38±10.52% vs 27.68±2.91% in *siControl*), while at the same time suppressing centriole disengagement (Fig. 8A-C). These results are consistent with a model in which Plk1 promotes premature centriole disengagement via a Separase-induced cleavage of the Pericentrin ring around centrioles.

**Figure 8:**
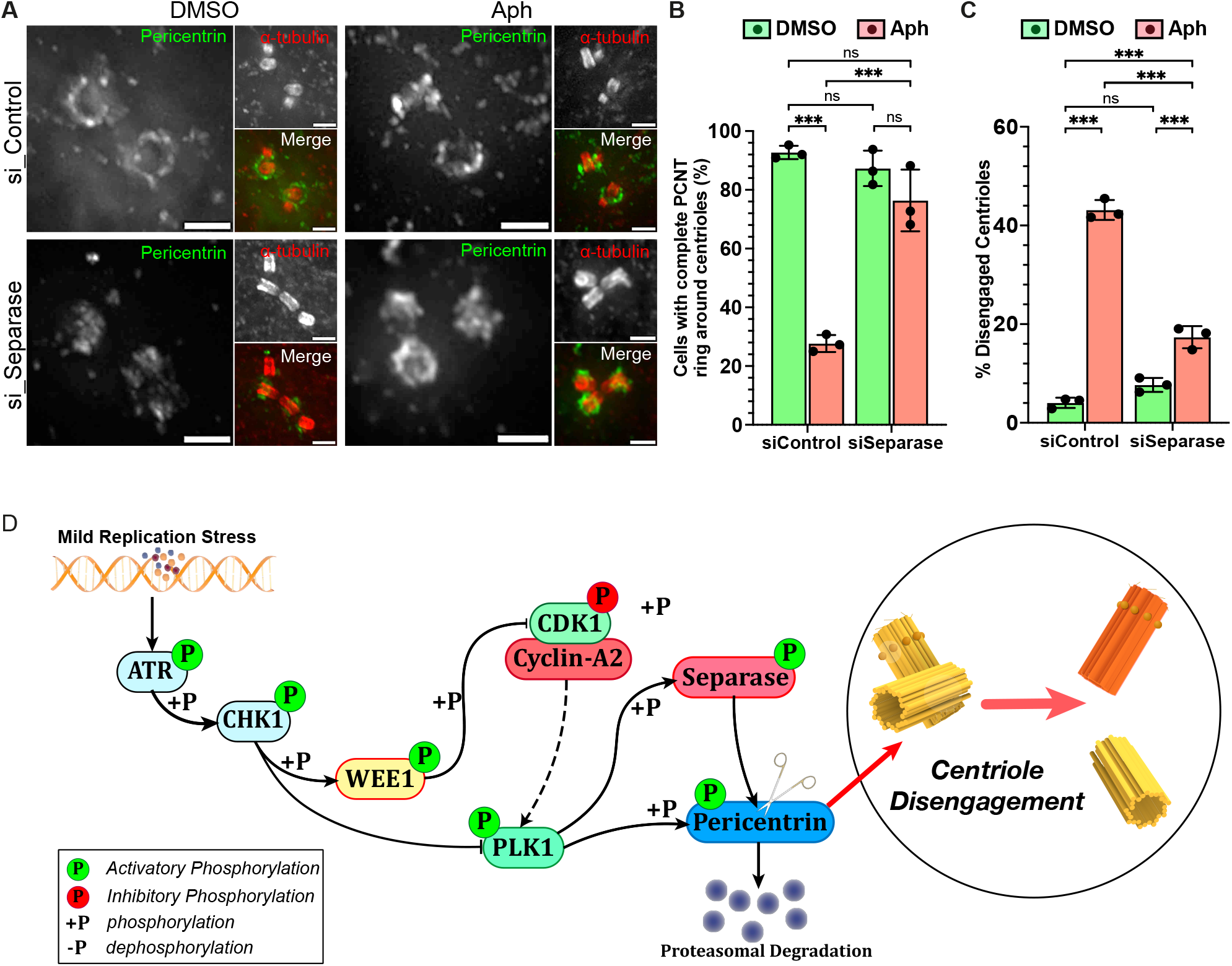
Premature centriole disengagement requires Separase dependent PCM disintegration. **(A)** Expansion microscopy images of centrioles in G2 phase hTERT-RPE1 cells treated either with DMSO or Aph and siControl or siSeparase and stained with α-tubulin (red) and Pericentrin (PCM: green). **(B)** Quantification of percentage of G2 phase cells with complete Pericentrin ring around their centrioles (*N* = 3 independent experiments, n = 68, 70, 60 and 61 cells treated with siControl:DMSO, siControl:Aph, siSeparase:DMSO and siSeparase:Aph, respectively). **(C)** Quantification of percentage of G2 phase cells with engaged centrioles in A (*N* = 3 independent experiments, n = 68, 70, 60 and 61 cells treated with siControl:DMSO, siControl:Aph, siSeparase:DMSO and siSeparase:Aph, respectively). Data presented as Mean values ± SD. (p > 0.05 = ns; p < 0.001 = ***: Sídak test). Scale bars = 0.5 µm. **(D)** Schematic representation of the molecular pathway regulating premature centriole disengagement during mild replication stress.

## Discussion

Here we investigated how mild replication stress in non-transformed cells disrupts the synchrony of the DNA and the centriole cycle, leading to premature centriole disengagement. Using expansion microscopy, we demonstrate that mild replication stress induces premature centriole disengagement already in G2 via the ATR-Chk1-Wee1 axis of the DNA damage repair pathway. Activation of this pathway dampens but does not block the activity of the mitotic kinase Plk1, a critical regulator of both the DNA and the centrosome cycle. A sub-critical Plk1 activity is insufficient to promote rapid mitotic entry, resulting in a G2 delay; it is, however, sufficient to drive centriole disengagement via Separase and the disassembly of the Pericentrin ring in the PCM, consistent with the canonical centriole disengagement pathway^8, 44^. We thus propose that the differential threshold in Plk1 activity for mitotic entry and centriole disengagement is at the origin of the asynchrony in DNA and centrosome cycle in cells experiencing mild replication stress.

In a previous study we had found that mild replication stress led to premature centriole disengagement and transient multipolar spindles in mitotic cells. When exactly centrioles became dis-engaged was nevertheless unknown, due to the insufficient resolution of classical fluorescence microscopy. Using expansion microscopy, we now demonstrate that mild replication stress already causes centriole disengagement in G2. Given that expansion microscopy allows to study a high number of cells per experiment, this method has the potential to replace electron microscopy as the gold standard to determine whether centriole pairs are engaged or not. Our results further indicate that mild replication stress induces centriole disengagement via the ATR-Chk1-Wee1 pathway, but not the ATM-Chk2 pathway. This indicates that under these conditions disengagement is controlled by the single-strand break (SSB) arm of the DNA damage repair pathway. Whether activation of the ATM-Chk2 double strand-break arm of the of DNA damage pathway also has the potential to create conditions favouring centriole disengagement remains to be seen.

The fact that centriole disengagement in G2 also depends on Cdk1/Cyclin A and Plk1, may at first appear counterintuitive since activation of the DNA damage repair pathway defers mitotic entry by preventing Cdk1 and Plk1 activation until DNA repair is completed^42, 45^. Nevertheless, here we find that mild replication stress does not fully inhibit Plk1 activity; rather our FRET measurements indicate that G2 cells with mild replication stress harbour a Plk1 activity around 60% when compared to unperturbed mitotic cells or when compared to G2 cells treated with Aphidicolin and an ATR inhibitor. We therefore postulate that mild replication stress reduces Plk1 activity via ATR to levels that are incompatible with a rapid mitotic entry, yet sufficient to drive centriole disengagement. Our data are consistent with a model in which both processes require different Plk1 activity thresholds, explaining how mild replication stress disrupts the synchronicity between the DNA and the centrosome cycle in late G2. Our model is also consistent with a previous study showing how severe DNA damage conditions that block mitotic entry can also induce a Plk1-dependent centriole disengagement in 40% of the cells after a long (48-72 hours) G2 arrest^46^. We speculate the lower kinetics of centriole disengagement under those conditions is due to lower residual Plk1 activity, consistent with the fact that the vast majority of the cells never enter mitosis.

If our hypothesis is correct, one key question is why a high Plk1 activity, which is present in unperturbed G2 cells, is not sufficient to drive centriole disengagement on its own. Under normal conditions centriole disengagement occurs at the end of mitosis when Cdk1/Cyclin-B1 has been inactivated by APC^Cdc20^ mediated protein degradation^47^ and Plk1 activity started to drop due to APC/C^Cdh1^ mediated degradation^48^. This could point to an inhibitory role of Cdk1/CyclinB1 during late G2 and mitosis that prevents a high Plk1 activity from inducing premature centriole disengagement. This could be achieved by the near simultaneous activation of both kinases during mitotic entry, which under normal conditions prevents an exclusive activation of Plk1. One plausible target of Cdk1/Cyclin B1 could be Separase, which we identify as a key regulatory of premature centriole disengagement, and which is inhibited by Cdk1/CyclinB1 activity^49^. The synchronous loss of Securin and Cyclin-B1 at anaphase onset by APC/C^Cdc20^ could thus not only activate Separase to induce chromosome segregation^50^, but also permit centriole disengagement under the control of Plk1. A second important question is how Separase can act on centrosomes in G2 in Aphidicolin-treated cells without affecting chromosome cohesion. We reason that one main reason is that Separase is excluded from the nucleus before mitotic entry by CRM1-dependent nuclear export to prevent cohesin cleavage during chromosome condensation^51^. Furthermore, cytosolic Separase is thought to be inactivated by Securin in addition to Cdk1 phosphorylation at S1126^52, 53^. Our results indicate that in the absence of robust Cdk1/Cyclin B1 activity, Securin inhibition is not sufficient, resulting in premature centriole premature centriole disengagement.

Downstream of Separase we identify PCM integrity and in particular loss of the structural integrity of the Pericentrin ring as likely pre-requirement for premature centriole disengagement in cells experiencing mild replication stress. This would indicate that premature centriole disengagement in G2 depends in part on the canonical centriole disengagement pathway in which Separase-induced cleavage at R2231 initiates Pericentrin degradation during mitotic exit^9^. We did, however, not observe any change in Cep57 localization, pointing to potential differences between the two processes. Whether premature centriole disengagement purely relies on Pericentrin cleavage or whether other components are involved remains an open question.

We conclude that replication stress in non-cancerous cells deregulates the synchrony between the cell and centriole duplication cycle and thus connects the DNA damage response pathway and the centriole disengagement via Plk1. This deregulation is much more pernicious than the premature centriole disengagement seen after strong DNA damage^46^, since only cells experiencing mild replication stress will nevertheless enter mitosis, forming transient multipolar spindles. Replication stress and deregulation of the centrosome cycle are both hallmarks of pre-cancerous and cancerous lesions^54-57^. To which extent the two phenomena are linked is, however, not clear. Our study uncovers a new connection between both processes, providing a possible mechanistic explanation for their co-evolution in cancer cells. Multiple theories do exist pertaining to the origin centrosome cycle deregulation and appearance of supernumerary centrosomes in cancer cells, including cytokinesis failure, uncontrolled centriole duplication or elongation and PCM disintegration during mitosis^58, 59^. Here, we speculate that a lack of synchrony between the centrosome and DNA cycle may also favour the appearance of abnormal centrosome number in cells experiencing persistent replication stress, as cells might accumulate dis-engaged centrioles that are prematurely licensed for centrosome duplication.

## Materials and Methods

### Cell culture and drug treatments

hTERT-RPE1 cells (ATCC: CRL-4000), hTERT-RPE1 EB3-eGFP/H2B-mCherry cells (kind gift by Willy Krek, ETH Zurich, Switzerland), hTERT-RPE1 OSTR1 cells, hTERT-RPE1 Cyclin-A2-double degron cells and hTERT-RPE1 Cyclin-B1-double degron cells (all kind gifts by Helfrid Hochegger, University of Sussex, United Kingdom) were cultured in Dulbecco’s Modified Eagle’s Medium (Thermo Fisher Scientific: 61965-026) supplemented with 10% foetal calf serum (FCS) (Thermo Fisher Scientific: 10902-096) and 100U/ml of each penicillin and streptomycin (P/S) (Thermo Fisher Scientific: 15140122) at 37°C, 95% relative humidity and 5% CO_2_ in humidified CO_2_ incubator. hTERT-RPE1 Plk1 FRET Sensor cells were prepared by transfecting Plk1-FRET sensor c-jun substrate plasmid^37^ (Addgene: 45203) in hTERT-RPE1 cells using X-tremeGENE™ 9 (Merck: XTG9-RO) transfection reagent according to manufacturer’s instructions. Transfected cells were selected with 600 µg/ml of G418 (Invivogen: ant-gn-5) in DMEM with 10% FCS and 100U/ml P/S followed by single cell cloning. All cell lines were routinely tested for mycoplasma contamination by PCR. For live-cell imaging, cells were cultured in Leibovitz’s L-15 medium without Phenol Red (Thermo Fischer Scientific: 21083-027) with 10% FCS and 100 U/ml P/S at 37°C.

To induce mild replication stress in all cell types Aphidicolin (Sigma Aldrich: A0781) was dissolved in DMSO and applied at 400 nM for 16 hrs. For auxin induced degradation of cyclins, cells were treated with 1 µg/ml Doxycycline (Sigma Aldrich: D9891-1G) for 2 hrs followed by addition of 3 µM Asunaprevir (Apexio: BMS-650032) and 500 µM 3-Indoleacetic acid (Sigma Aldrich: I2886). Cells were treated with following inhibitors to inhibit indicated kinases, Cdk1: 9 µM RO3306 (Sigma Aldrich: SML0569), Plk1: 10 nM BI2536 (Selleck Chemicals: S1109), ATR: 0.8 µM ETP-46464 (Apexbio: A8626), ATM: 10 µM KU-55933 (Selleck Chemicals: S1092), Chk1: 5 µM LY2603618 (Chk1i-1) (Selleck Chemicals: S2626), Chk1: 1 µM PF-477736 (Selleck Chemicals: S2904), Chk2: 10 µM BML-277 (Selleck Chemicals: S8632), Wee1: 0.5 µM PD0166285 (Wee1i-1) (Selleck Chemicals: S8148), and Wee1: 0.5 µM MK-1775 (Selleck Chemicals: S1525).

### RNA Interference

siRNA transfections were performed using Lipofectamine RNAiMAX (Thermo Fisher Scientific: 13778075) according to manufacturer’s instructions. RNAi was performed for 72 h; when combined with inhibitors the drugs were added 60 hours post transfection. All the siRNA sequences used in the study were previously validated sequences: siControl (Qiagen, GGACCTGGAGGTCTGCTGT) and siSeparase (Dharmacon, GCTTGTGATGTCATGCCATCCTGA)^60^.

### Antibodies

The following antibodies were used in this study: Mouse anti-α-tubulin (Geneva antibody facility: AA345-M2a; 1:250: ExM)^61^, Mouse anti-β-tubulin (Geneva antibody facility: AA344-M2a; 1:250: ExM)^61^, Mouse anti-α-tubulin (Sigma Aldrich, T9026, 1:5000: Western Blotting), Rabbit anti-Pericentrin (abcam: ab4448; 1:250: ExM, 1:1000: STED), Rabbit anti-CEP57 (GeneTex: GTX115931; 1:250: ExM), Mouse anti-Cyclin-A2 (abcam: ab38; 1:1000: Western Blotting), Mouse anti-Cyclin-B1 (abcam: ab72; 1:1000: Western Blotting) and Mouse anti-CENP-F (abcam: ab90; 1:1000: STED). All the Alexa Fluor-conjugated secondary antibodies were purchased from Thermo Fisher Scientific and used at 1:500 dilution. HRP-Conjugated goat anti-mouse antibody for western blotting was purchased from Thermo Fisher (Cat # 32430) and used at 1:10,000 dilution. Goat anti-rabbit STAR RED antibody for STED microscopy, purchased from Abberior instruments GmbH (Cat # 2-0002-011-9) was used at 1:1000 dilution.

### Expansion microscopy

Cells were grown on 12 mm circular glass coverslips (Thermo Fisher Scientific) and treated with required inhibitors/drugs overnight. Next day the coverslips were treated with Acrylamide (AA)-Formaldehyde (FA) Solution [1.4% AA (Sigma Aldrich: A4058) and 2% FA (Sigma Aldrich: F8775) in PBS] for 5 h at 37°C to prevent protein crosslinking. Coverslips were next subjected to gelation by incubation for 1 hr at 37°C with monomer solution [19% Sodium Acrylate (Sigma Aldrich: 408220), 10% Acrylamide, 0.1% Bisacrylamide (Sigma Aldrich: M1533), 0.5% Tetramethyl ethylenediamine-TEMED (Thermo Fisher: 17919), 0.5% Ammonium Persulfate (Thermo Fisher: 17874) in PBS]. Post gelation, the coverslips were treated with denaturation solution [50 mM Tris (Sigma Aldrich: 99362), 200 mM Sodium dodecyl sulphate (Axon Lab AG: A2572.0500), 200 mM Sodium Chloride (Axon Lab AG: A3597.1000) in Nuclease free water, pH: 9.0] for 15 min on a rocker shaker at room temperature to detach the gels from coverslips. The gels were heated at 95°C for 90 min in denaturation solution followed by three 30 min washes with water. The gels were incubated with PBS for 15 min followed by 3 hrs incubations with primary and secondary antibodies followed each by three 10 min washes at 37°C and gentle shaking. Stained gels were kept overnight in water for optimal expansion. The size of gel was measured to calculate the expansion factor and the gel was cut into small pieces and placed in 2 well plastic bottom ibidi chamber (Ibidi GMBH: Cat # 80286)- The 3D image stacks of centrioles in G2 phase cells (4 centrioles in one cell) were acquired in 0.1 µm steps using a 100x oil-immersion (NA 1.4) objective on an Olympus DeltaVision microscope (GE Healthcare) equipped with a DAPI/FITC/Rhodamine/CY5 filter set (Chroma Technology Corp) and a CoolSNAP HQ camera (Roper-Scientific). The three-dimensional image stacks were deconvolved with SoftWorx (GE Healthcare). The acquired images were cropped and processed with imageJ (NIH) software to construct 3D image to analyse the configuration of centrioles (orthogonal orientation and distance in between) for each image.

### Live cell imaging and analysis

For live cell imaging experiments hTERT-RPE1 cells stably expressing EB3-eGFP and H2B-mCherry were plated in glass bottom Ibidi chambers (Ibidi GMBH: Cat # 81158) and normal DMEM medium was replaced with L15 Leibovitz’s medium supplemented with 10% FCS and 100 U/ml P/S. The cells were treated with indicated inhibitors and imaged at 37 °C on a Nikon Ti microscope equipped with a 60x NA 1.3 oil objective, DAPI/FITC/Rhodamine/CY5 (Chroma, USA) filter set, Orca Flash 4.0 CMOS camera (Hamamatsu, Japan) and the NIS software. Cells were recorded every 3 min for 18 h with z-slices separated by 2 μm, and 100 ms exposure per z-slice at wavelengths of 488 (525) and 561 nm (615 nm) excitation (emission). The time-lapse movies were analysed manually for multipolarity using Imaris software (Bitplane Inc).

### STED nanoscopy

Cells were grown on glass coverslips (170±10 µm thick-Hecht-Assistent: Cat # 41014515) and fixed with chilled methanol at −20°C for 6 min. Coverslips were stained with Rabbit anti Pericentrin and Mouse anti-CENP-F (to identify G2 phase cells). Coverslips with fluorescent labelled samples were mounted on microscope slides using Fluoromount-G™ (Thermo Fisher Scientific: Cat # 00-4958-02) and sealed with nail paint from all sides. Dual colour 2D-STED imaging was performed on a TCS SP8 STED microscope (Leica Microsystems) at 21°C using a STED motorized oil immersion objective (HCPL Apo 100×/NA 1.30 motCOR) using LAS-X Imaging software (Leica Microsystems). Excitation was performed with a white light laser (WLL), and depletion with a 775-nm pulsed laser. Both the excitation and depletion lasers were calibrated either with the STED Expert Alignment Mode and Abberior gold nanoparticles (diameter: 80 nm) before starting each imaging session, or with the STED Auto Beam Alignment tool during imaging sessions (Leica LAS X software). The STED imaging was made sequentially using excitation at 580 nm (WLL) and a STED 775 depletion laser line for Abberior STAR RED (for Pericentrin) anti-rabbit antibody. Detection signals were collected between 647 and 677 nm for STAR RED using highly sensitive Leica Hybrid Detectors with a fixed gain and offset (100 mV and 0, respectively). Time*-*gated detection was used for all fluorophores (0.50–6.00 ns). Acquisitions were performed with a line average of 4, a speed of 400 Hz, and software optimized pixel size respecting the Nyquist criteria. 2D-STED images were deconvolved using the Leica Lightning Mode (LAS X software) and the analysis for Pericentrin structure was performed with ImageJ (National Institutes of Health). Cells were co-stained with CENP-F in AF488 channel and the signal was used only to identify late G2 phase cells.

### FRET assay to measure Plk1 activity

hTERT-RPE1 Cells stably expressing Plk1 FRET sensor were seeded in four-well glass bottom µ-Slide ibidi chambers (Ibidi; 80426) and treated with different inhibitors/drugs as indicated in DMEM with 10% FCS and 1% P/S. The DMEM was replaced with Leibovitz L-15 supplemented with 10% FCS and 1% P/S containing same drugs/inhibitors as before. The chambers were acclimatised in 37 °C chamber before imaging. The acquisition was performed with an EC Plan Neofluor 100X (NA 1.3) oil objective on a Zeiss Cell Observer.Z1 spinning disk microscope (Nipkow Disk) equipped with a 37 °C chamber and a CSU X1 automatic Yokogowa spinning disk head. To perform FRET experiments samples were illuminated with 445 nm laser and the emission signal was split equally using DV2 split view system and CFP and YFP emissions were recorded on the split beams. 512 ×512 pixel size images were acquired with an Evolve EM512 camera (Photometrics) using Visiview 4.00.10 software. Acquired images were analysed using ImageJ to calculate the YFP to CFP emission intensity ratio after background subtraction.

### Immunoblotting

Cells were grown in 60 mm plastic dishes and treated with inhibitors/drugs overnight. To prepare protein lysate the cells were scrapped off using cell scrapper and lysed in RIPA buffer (50 mM Tris pH-7.4, 150 mM NaCl 1% Nonidet P-40 (Thermo Fisher Scientific: 85124), 0.5% Sodium deoxycholate (Sigma Aldrich: D5670), 0.1% Sodium dodecyl-sulphate in ultrapure water) supplemented with Protease inhibitor (Roche: 11873580001) and Phospho-STOP (Roche: 04906845001). Protein concentrations in the lysates were determined using the Bradford Protein Assay (Thermo Fisher; 23200). Samples with equal amounts of protein were mixed with 5X Laemmli buffer and heated to 95 °C for 5 min. Proteins were separated on a 10% SDS-polyacrylamide gels and transferred onto a 0.45 µm pore size nitrocellulose membrane (Macherey-Nagel GMBH: 741280) by wet blotting. Membranes were blocked with 5% non-fat dry milk in PBS 0.2% Tween20 (PBS-T) for 30 min. After blocking, membranes were incubated with primary antibodies overnight at 4 °C in PBS-T 5% non-fat dry milk. Membranes were washed three times with PBS-T and incubated 1 h with the appropriate peroxidase-conjugated secondary antibody in PBS-T 5% non-fat dry milk. The membranes were washed thrice with PBST and the bands corresponding to protein of interest were detected by chemiluminescence using the Amersham ECL Prime Western Blotting Detection Kit (GE Healthcare; RPN2232) in a Fusion FX7 Spectra Multispectral Imaging system (Witec AG, Switzerland).

### Statistical analysis

Statistical tests for all figures were performed using GraphPad Prism 9 (GraphPad), the statistical tests employed in every case are described in the figure legends. Minimum three independent biological replicates were performed in all experiments.

## Acknowledgements

Authors thank H. Hochegger (University of Sussex, UK) for cell lines, P. Guichard, M. Laporte and V. Hamel (University of Geneva, Switzerland) for expansion microscopy support, Members of Bioimaging and FACS facility (University of Geneva, Switzerland) for experimental support, Monica Gotta and team (University of Geneva, Switzerland) and members of Meraldi laboratory for helpful discussions and support, and P. Guichard, V. Hamel and M. Gotta for critical comments on the manuscript.

## Competing interests

The authors declare no competing or financial interests.

## Author contributions

Conceptualization: D.D., P.M.; Formal analysis: D.D., P.M.; Investigation: D.D., D.H.; Writing - original draft: D.D.; Writing - review & editing: D.D., P.M.; Visualization: D.D.; Supervision: P.M.; Project administration: P.M.; Funding acquisition: P.M.

## Funding

This work in Meraldi Lab was supported by the Swiss National Science Foundation (Schweizerischer Nationalfonds zur Förderung der Wissenschaftlichen Forschung; SNF) project grant (No. 31003A_179413) and the Université de Genève.

## Figure Legends

**Figure S1:**
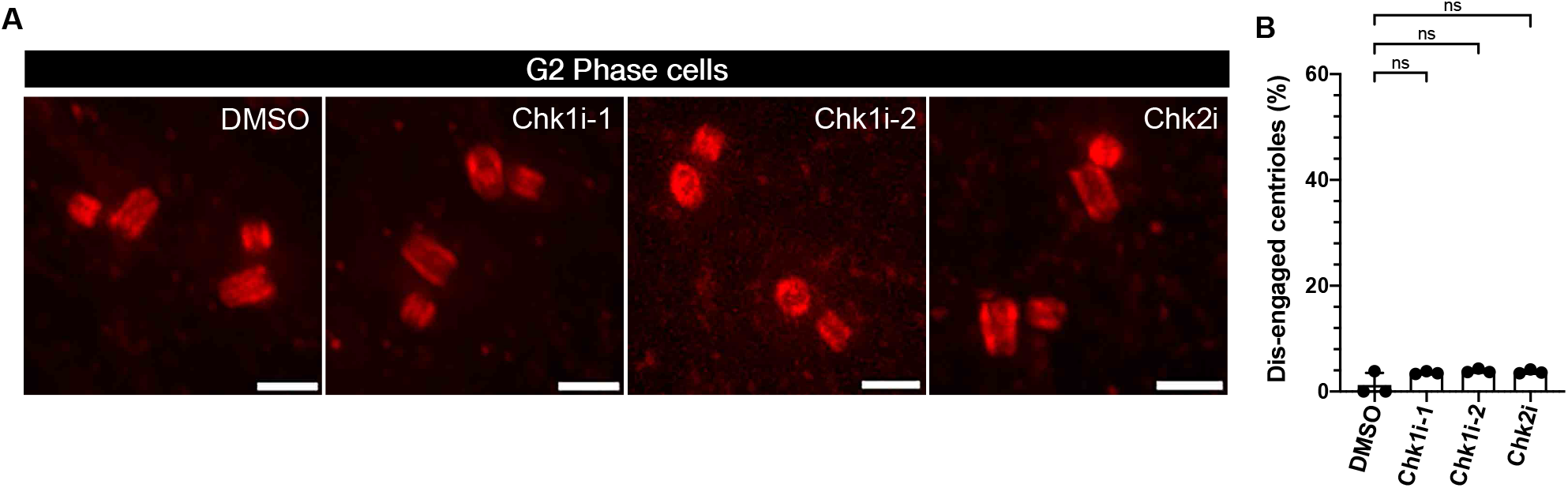
Inhibition of Chk1 or Chk2 alone did not cause premature centriole disengagement in G2 phase: **(A)** Expansion microscopy images of centrioles, stained with α-tubulin, in G2 phase hTERT-RPE1 cells treated with indicated drugs/inhibitors. **(B)** Quantification of percentage of G2 phase cells (shown in A) having engaged centrioles within their centrosomes (*N* = 3 independent experiments, n = 78, 85, 77 and 81 cells in DMSO, Chk1i-1, Chk1i-2 and Chk2i, respectively) Data presented as Mean values ± SD. (p > 0.05 = ns; Sídak test). Scale bars = 0.5 µm.

**Figure S2:**
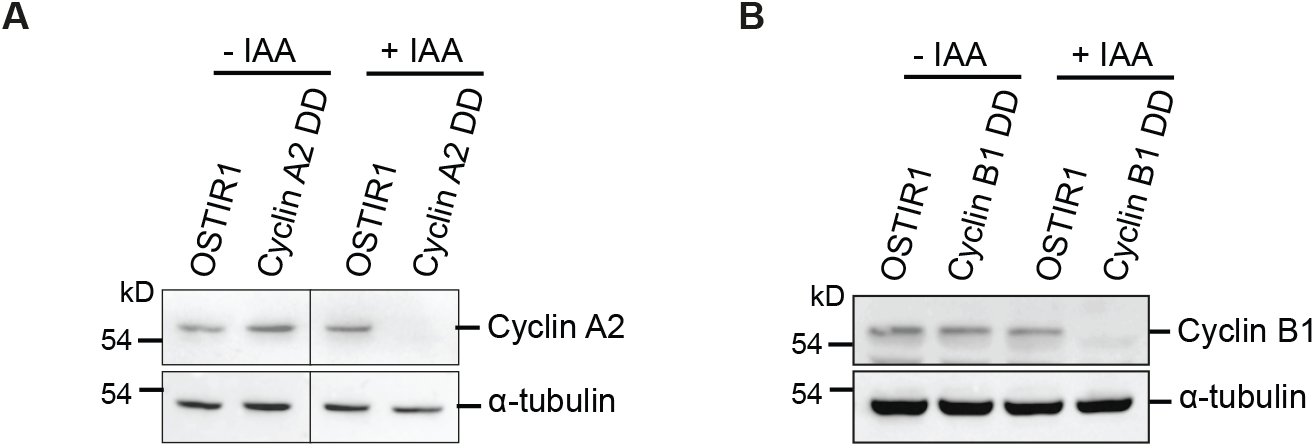
Auxin treatment causes degradation of cyclins: **(A)** Western blot image showing degradation of Cyclin-A2 in Cyclin-A2 double degron containing RPE1 cells after addition of Auxin (IAA) but not in OSTIR Control RPE1 cells. The blots were also probed with α-tubulin to confirm equal protein loading. **(B)** Western blot image showing degradation of Cyclin-B1 in Cyclin-B1 double degron containing RPE1 cells after addition of Auxin (IAA) but not in OSTIR Control RPE1 cells. The blots were also probed with α-tubulin to confirm equal protein loading.

**Figure S3:**
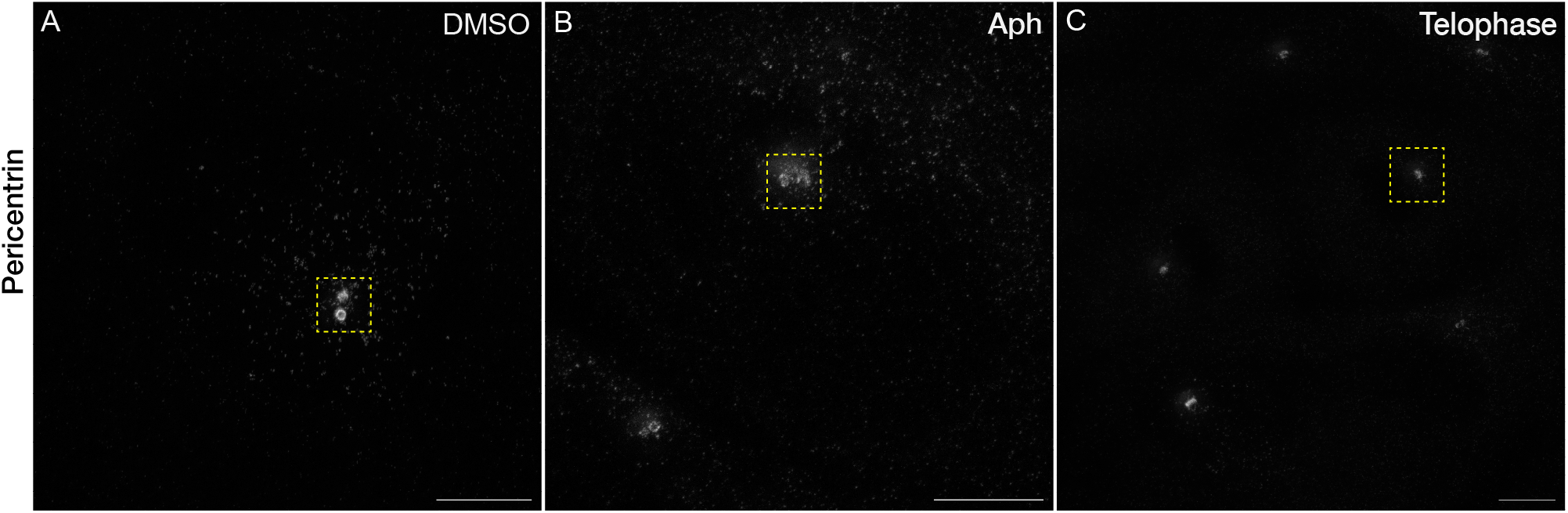
Replication Stress disrupts PCM integrity: **(A-C)** STED nanoscopy images of G2 phase hTERT-RPE1 cells treated either with DMSO **(A)** or Aph **(B)** and DMSO treated cells in telophase **(C)**. The cells were stained for CENP-F for identification of G2 cells. Insets represent the pericentriolar region shown below and in Fig 7A (in grey). Scale bars = 5 µm.

### Movies

**Movie 1:** Live-cell imaging of hTERT-RPE1 stably expressing EB3-GFP and H2B-mCherry and treated with DMSO control. Every image was captured at an interval of 3 m. The movie is shown at 5 frames per second (fps). Scale bars, 5 µm.

**Movie 2:** Live-cell imaging of hTERT-RPE1 stably expressing EB3-GFP and H2B-mCherry and treated with Aphidicolin. Every image was captured at an interval of 3 m. The movie is shown at 5 frames per second (fps). Scale bars, 5 µm.

**Movie 3:** Live-cell imaging of hTERT-RPE1 stably expressing EB3-GFP and H2B-mCherry and treated with Wee1i-1. Every image was captured at an interval of 3 m. The movie is shown at 5 frames per second (fps). Scale bars, 5 µm.

**Movie 4:** Live-cell imaging of hTERT-RPE1 stably expressing EB3-GFP and H2B-mCherry and treated with Wee1i-2. Every image was captured at an interval of 3 m. The movie is shown at 5 frames per second (fps). Scale bars, 5 µm.

**Movie 5:** Live-cell imaging of hTERT-RPE1 stably expressing EB3-GFP and H2B-mCherry and treated with Aphidicolin and Wee1i-1. Every image was captured at an interval of 3 m. The movie is shown at 5 frames per second (fps). Scale bars, 5 µm.

**Movie 6:** Live-cell imaging of hTERT-RPE1 stably expressing EB3-GFP and H2B-mCherry and treated with Aphidicolin and Wee1i-2. Every image was captured at an interval of 3 m. The movie is shown at 5 frames per second (fps). Scale bars, 5 µm.

## Notes

### Competing Interest Statement

The authors have declared no competing interest.

